# Digit Ratio (2D:4D) and Amniotic Testosterone and Estradiol: An Attempted Replication of Lutchmaya et al. (2004)

**DOI:** 10.1101/2020.07.10.197269

**Authors:** Gareth Richards, Wendy V. Browne, Mihaela Constantinescu

## Abstract

The ratio of length between the second (index) and fourth (ring) fingers (digit ratio or 2D:4D) is frequently employed as a retrospective marker of prenatal sex hormone exposure. Lutchmaya et al. (2004) reported that the ratio of testosterone (T) to estradiol (E) present in second trimester amniotic fluid was negatively correlated with digit ratios for the right hand (but not the left hand) in a sample of 29 children at 2-year follow-up. This observation is frequently cited as evidence for the measure’s validity but has not been replicated. We therefore present the findings of another study of amniotic T and E that did not find evidence for these effects at 4½-year follow-up. The confidence intervals were large, the direction of correlations observed was generally erratic, and the overall findings therefore question the premise that second trimester sex hormones affect the development of digit length ratios in humans.

## 1. Introduction

Manning et al.^1^ suggested that the ratio of length between the second and fourth fingers (digit ratio or 2D:4D) is a negative correlate of prenatal testosterone exposure and a positive correlate of prenatal oestrogen exposure. As there are considerable practical and ethical constraints to measuring prenatal hormones more directly, there has been much interest in utilising 2D:4D as a tool for retrospective examination of the developmental origins of sexually differentiated outcomes; however, its validity has frequently been questioned^2–4^.

Experimental manipulation of foetal sex hormones is not permitted in human studies for obvious ethical reasons, and so researchers have developed a range of creative approaches to address this problem. These include investigation of patient groups exposed to atypical sex hormone concentrations (or sensitivity), such as congenital adrenal hyperplasia^5^ and androgen insensitivity syndrome^6,7^, and comparing same-sex and opposite-sex twins^8–10^. Although theory-consistent effects have been reported in a number of studies and across different methodologies, these are typically present alongside null findings and replication failures.

A more direct approach has been to measure sex hormone concentrations in amniotic fluid. Amniocentesis is an invasive medical procedure by which amniotic fluid is extracted for genetic and chromosomal analysis in at-risk pregnancies. Although such samples may not be representative of the general population, amniocentesis has routinely been performed in typically-developing pregnancies of advanced maternal age. Manning^11^ (see Figure 2.4, p. 32) initially reported that maternal 2D:4D was negatively correlated with the level of testosterone present in the amniotic fluid, although it has been questioned whether this association could have been inflated by the presence of outliers^12^. Lutchmaya et al.^13^ then reported that the ratio of testosterone to estradiol (T:E) present in amniotic fluid was significantly negatively correlated with R2D:4D in 29 two-year-old children. This finding indicates that a high level of testosterone relative to estradiol is associated with the development of a low, more ‘male-typical’, 2D:4D ratio in the right hand. However, the sample size was small, males and females were not analysed separately, no significant effect was observed for L2D:4D, and neither testosterone nor estradiol on its own was a significant predictor.

Although the study by Lutchmaya et al.^13^ is frequently cited in support of the validity of 2D:4D, in the 16 years since its inception, no direct replication attempt has been published. The closest has been a study reporting a significant negative correlation between amniotic testosterone and L2D:4D in female neonates^14^; however, no significant effect was observed for R2D:4D in females, or for R2D:4D or L2D:4D in males. A re-analysis of these data^4^ showed a significant negative correlation with the average of R2D:4D and L2D:4D (M2D:4D) in females (but not in males); notably, there was no correlation with the right-left difference in 2D:4D (D_[R-L]_), an additional variable for which low values have been hypothesised to reflect high levels of foetal androgen exposure^11^. Importantly, amniotic estradiol was not measured in this study, meaning that no attempt at replicating the significant effect reported by Lutchmaya et al.^13^ could be made.

The current paper reports the findings of a study in which we examined whether amniotic T, E, and T:E ratio were predictive of digit ratio variables measured in the children and mothers of these pregnancies at 4½ year follow-up.

## 2. Method

### 2.1 Participants

The sample for this study was obtained from women undergoing amniocentesis at the Queen Charlotte’s and Chelsea Hospital, London. Although the reason for amniocentesis was usually increased risk of Down syndrome, only women carrying healthy foetuses were retained. There were in total 66 mothers (with ages at birth ranging from 28.17 to 44.08 years; *M* = 37.68, *SD* = 4.01) from whom amniotic fluid samples were collected usually between weeks 15 and 22 of gestation. The women gave birth to 66 children (32 females, 34 males) whose digit ratios (along with those of their mothers) were measured around the age of 4½ years (range = 3.83–5.92, *M* = 4.501, *SD* = 0.629). Most of the women (78.8%, n=52) were Caucasian, 9.1% (n=6) were Asian, 6.1% (n=4) were African, 4.5% (n=3) were Middle-Eastern, and n=1 (1.5%) was of mixed ethnicity. Regarding education, 19.7% (n=13) had a postgraduate degree 43.9% (n=29) had an undergraduate degree, 16.7% (n=11) had vocational training, 9.1% (n=6) had A-levels, and 10.6% (n=7) had GCSEs or equivalent. Study procedures were approved by national and institutional research ethics committees and conducted in accordance with the Declaration of Helsinki, and written consent was obtained from all the mothers who took part in this research.

### 2.2 Amniotic Hormones

Amniotic fluid samples were obtained between 2002 and 2004 when women were recruited to the study as part of an ongoing larger-scale project examining associations between hormones and behaviour (see Bergman et al.^15^) Total testosterone (T) concentrations were measured in amniotic fluid samples by radioimmunoassay (RIA), Coat-a-Count (DPC Los Angeles, CA), with intra- and inter-assay coefficients of variation of 7.5% and 8.9%. A random subset (18 males; 12 females) of the samples was also analysed for T and E by Liquid Chromatography/Mass Spectroscopy (LCMS). The correlation between T values using RIA and LCMS was strong, *r*(40) = 0.82, *p* < 0.001^15^.

### 2.4 Digit ratio (2D:4D)

2D:4D data were collected between 2005 and 2009 at approximately 4½ years follow-up. Measurements were taken directly (from the hands) and/or indirectly (from photocopies) using callipers measuring to 0.01mm. The intra-class correlation (single measures, absolute agreement) for a subsample (n=15) of participants’ photocopies for R2D:4D determined that the repeatability of measurement was high, *ICC* = 0.940, *p* < 0.001. To maximise the sample size that could be included in the analysis (and thereby increasing statistical power), we used the following calculation^16^ to correct the measurements taken directly for those participants whose digit ratios had not also been measured from photocopies:

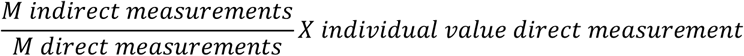

### 2.5 Statistical analysis

We used between subjects t tests to examine sex differences and Pearson’s correlations to test for associations between hormonal and digit ratio variables. We also calculated the bias corrected accelerated 95% confidence intervals (BCa 95% CI) based on 2,000 resamples. We took this approach because some variables were not normally distributed, and outliers were present for some of the hormonal variables. Using bootstrapping therefore allowed us to retain all biologically relevant data, and produce more reliable estimates than would be obtained from standard parametric analyses.

## 3. Results

Amniotic T (both RIA and LCMS measurements) and T:E ratio were significantly higher when the foetus was male, though there was no sex difference for E. There were no sex differences for R2D:4D, L2D:4D, and M2D:4D. D_[R-L]_ was marginally lower in males (*p* = 0.049), though the BCa 95% CIs overlapped zero (bootstrapped *p* = 0.057) (see **Table 1**).

**Table 1.** Sex differences for amniotic hormone and digit ratio variables.

|  | Males |  |  | Females |  |  | Difference |  |  |  |  |
| --- | --- | --- | --- | --- | --- | --- | --- | --- | --- | --- | --- |
|  | *n* | *M* | *SD* | *n* | *M* | *SD* | *t* | *p* | *d* | Mean dif. | BCa 95% CI |
| **T RIA (nmol/L)** | 34 | 0.858 | 0.441 | 31 | 0.256 | 0.157 | **-7.456<sub>§</sub>** | **< 0.001** | **-1.786** | **-0.602** | **-0.771, -0.453** |
| **T LCMS (pg/ml)** | 18 | 204.000 | 86.518 | 12 | 27.917 | 11.269 | **-8.527<sub>§</sub>** | **< 0.001** | **-2.598** | **-176.083** | **-218.439, -134.922** |
| **E (pg/ml)** | 18 | 70.000 | 86.979 | 12 | 56.917 | 31.782 | -0.583<sub>§</sub> | 0.566 | -0.185 | -13.083 | -59.472, 25.928 |
| **T:E (LCMS)** | 18 | 7.277 | 6.839 | 12 | 0.655 | 0.460 | **-4.094<sub>§</sub>** | **0.001** | **-1.241** | **-6.622** | **-10.100, -3.420** |
| **R2D:4D** | 34 | 0.940 | 0.030 | 32 | 0.944 | 0.042 | 0.368 | 0.714 | 0.110 | 0.004 | -0.015, 0.022 |
| **L2D:4D** | 34 | 0.951 | 0.033 | 32 | 0.938 | 0.040 | -1.484 | 0.143 | -0.356 | -0.013 | -0.030, 0.004 |
| **M2D:4D** | 34 | 0.946 | 0.028 | 32 | 0.941 | 0.036 | -0.627<sub>§</sub> | 0.533 | -0.156 | -0.005 | -0.022, 0.011 |
| **D<sub>[R-L]</sub>** | 34 | -0.011 | 0.029 | 32 | 0.006 | 0.038 | 2.010 | 0.049 | 0.505 | 0.017 | -0.0004, 0.035 |

*Note.* BCa 95% CI = bias corrected accelerated 95% confidence intervals (calculated for mean difference via the bootstrapping procedure); E = estradiol; LCMS = Liquid Chromatography/Mass Spectroscopy RIA = radioimmunoassay; T = testosterone; § equal variances not assumed.

Pearson’s tests with BCa (i.e. bootstrapped) 95% confidence intervals for associations between amniotic fluid sex hormone concentrations and both maternal and child digit ratio variables at 4½-year follow-up are shown in **Table 2**. LCMS T levels in females correlated positively with R2D:4D, and there was some limited evidence of there being similar effects for M2D:4D (BCa 95% CIs did not cross zero but the parametric analysis was only marginally significant, *p* = 0.054) and D_[R-L]_ (parametric analysis was significant, *p* = 0.030, but the BCa 95% CIs crossed zero). There was also a negative correlation between amniotic E and D_[R-L]_ in males (BCa 95% CIs did not cross zero but the parametric statistic was not significant, *p* = 0.060). Each of these effects was in the opposite direction to that which would be predicted by theory. The only other finding of note was that the T:E ratio in males correlated negatively with maternal L2D:4D; although this effect was in the theory-consistent direction, and the BCa 95% CI did not cross zero, the conventional parametric statistical test was not significant (*p* = 0.126). There were no statistically significant correlations between amniotic T, E, or T:E ratio and any of the digit ratio variables measured in the children at follow-up.

**Table 2.** Associations between amniotic sex hormone concentrations and children’s digit ratio variables.

|  | Sex | 2D:4D | R2D:4D |  |  |  | L2D:4D |  |  |  | M2D:4D |  |  |  | D[R-L] |  |  |  |
| --- | --- | --- | --- | --- | --- | --- | --- | --- | --- | --- | --- | --- | --- | --- | --- | --- | --- | --- |
|  |  |  | *n* | *r* | *p* | *BCa 95% CI* | *n* | *r* | *p* | *BCa 95% CI* | *n* | *r* | *p* | *BCa 95% CI* | *n* | *r* | *p* | *BCa 95% CI* |
| **RIA T** | Males | Child | 34 | -0.065 | 0.713 | -0.335, 0.219 | 34 | 0.058 | 0.746 | -0.185, 0.310 | 34 | -0.002 | 0.993 | -0.236, 0.249 | 34 | -0.135 | 0.447 | -0.383, 0.089 |
|  |  | Mother | 32 | 0.059 | 0.749 | -0.191, 0.283 | 32 | -0.060 | 0.742 | -0.312, 0.231 | 32 | 0.004 | 0.984 | -0.234, 0.239 | 32 | 0.126 | 0.491 | -0.191, 0.465 |
|  | Females | Child | 31 | 0.069 | 0.711 | -0.239, 0.333 | 31 | -0.019 | 0.919 | -0.326, 0.255 | 31 | 0.027 | 0.883 | -0.287, 0.275 | 31 | 0.091 | 0.625 | -0.257, 0.441 |
|  |  | Mother | 30 | -0.067 | 0.724 | -0.316, 0.218 | 30 | -0.007 | 0.969 | -0.321, 0.301 | 30 | -0.046 | 0.809 | -0.298, 0.220 | 30 | -0.080 | 0.674 | -0.318, 0.133 |
| **LCMS T** | Males | Child | 18 | -0.262 | 0.293 | -0.598, 0.072 | 18 | 0.031 | 0.902 | -0.369, 0.499 | 18 | -0.130 | 0.607 | -0.446, 0.238 | 18 | -0.367 | 0.134 | -0.705, 0.063 |
|  |  | Mother | 16 | -0.059 | 0.827 | -0.570, 0.662 | 16 | 0.159 | 0.556 | -0.319, 0.627 | 16 | 0.056 | 0.837 | -0.379, 0.525 | 16 | -0.215 | 0.424 | -0.698, 0.494 |
|  | Females | Child | 12 | 0.466 | 0.127 | -0.296, 0.865 | 12 | 0.252 | 0.429 | -0.251, 0.681 | 12 | 0.426 | 0.168 | -0.145, 0.831 | 12 | 0.228 | 0.477 | -0.405, 0.663 |
|  |  | Mother | 12 | 0.655 | **0.021** | **0.053, 0.891** | 12 | 0.382 | 0.220 | -0.171, 0.776 | 12 | 0.567 | 0.054 | **0.112, 0.814** | 12 | 0.623 | **0.030** | -0.168, 0.899 |
| **Amniotic E** | Males | Child | 18 | -0.217 | 0.387 | -0.482, 0.125 | 18 | -0.080 | 0.752 | -0.276, 0.184 | 18 | -0.164 | 0.514 | -0.435, 0.256 | 18 | -0.177 | 0.483 | -0.579, 0.289 |
|  |  | Mother | 16 | -0.097 | 0.720 | -0.611, 0.337 | 16 | 0.392 | 0.134 | -0.290, 0.839 | 16 | 0.166 | 0.538 | -0.451, 0.667 | 16 | -0.481 | 0.060 | **-0.741, -0.122** |
|  | Females | Child | 12 | 0.056 | 0.862 | -0.595, 0.591 | 12 | -0.061 | 0.849 | -0.480, 0.338 | 12 | -0.001 | 0.998 | -0.613, 0.582 | 12 | 0.112 | 0.729 | -0.383, 0.594 |
|  |  | Mother | 12 | 0.155 | 0.631 | -0.603, 0.831 | 12 | 0.121 | 0.708 | -0.399, 0.712 | 12 | 0.148 | 0.647 | -0.425, 0.754 | 12 | 0.109 | 0.735 | -0.750, 0.757 |
| **Amniotic T:E** | Males | Child | 18 | -0.275 | 0.269 | -0.664, 0.288 | 18 | 0.035 | 0.890 | -0.254, 0.488 | 18 | -0.135 | 0.592 | -0.467, 0.268 | 18 | -0.388 | 0.111 | -0.729, 0.258 |
|  |  | Mother | 16 | 0.097 | 0.720 | -0.566, 0.626 | 16 | -0.399 | 0.126 | **-0.691, -0.121** | 16 | -0.171 | 0.527 | -0.638, 0.279 | 16 | 0.488 | 0.055 | -0.075, 0.850 |
|  | Females | Child | 12 | 0.298 | 0.347 | -0.231, 0.807 | 12 | 0.300 | 0.344 | -0.225, 0.646 | 12 | 0.351 | 0.264 | -0.139, 0.632 | 12 | 0.017 | 0.957 | -0.423, 0.432 |
|  |  | Mother | 12 | 0.059 | 0.855 | -0.511, 0.512 | 12 | 0.038 | 0.906 | -0.426, 0.376 | 12 | 0.053 | 0.870 | -0.444, 0.429 | 12 | 0.052 | 0.873 | -0.619, 0.631 |

*Note.* *BCa 95% CI* = bias corrected accelerated 95% confidence intervals (calculated via the bootstrapping procedure); *E* = estradiol; *LCMS* = Liquid Chromatography/Mass Spectroscopy; *RIA* = radioimmunoassay; *T* = testosterone; all statistical tests were Pearson's correlations (two-tailed)

## 4. Discussion

We report the findings of a study examining associations between individual differences in amniotic sex hormone concentrations and digit ratio (2D:4D). Although previous reports have suggested that high levels of amniotic T are associated with low digit ratios in both mothers^11^ and neonates^14^, and that high ratios of amniotic T:E are associated with low digit ratios in two-year-old infants^13^, we did not find evidence for such effects here. The findings therefore cast doubt on the idea that mid-trimester sex hormone concentrations are instrumental in the development of 2D:4D ratios in humans.

Most correlations between amniotic hormone levels and maternal digit ratio variables were not statistically significant, and the direction of these correlations was generally erratic. Of the five correlations for which some degree of statistical significance was indicated (i.e. a parametric *p* < 0.05 and/or BCa 95% CIs that did not cross zero), only one was in the theory-consistent direction. This was a negative correlation between T:E ratio in female pregnancies and the maternal L2D:4D. However, this effect should be interpreted with considerable caution considering (i) the high number of statistical tests that were run (and that we did not adjust for alpha inflation), (ii) the small sample size (n=12) for this analysis, and (iii) that although the BCa 95% CIs did not cross zero the conventional parametric statistical test was not significant (*p* = 0.126), and (iv) that similar effects were not observed for the other digit ratio variables (i.e. R2D:4D, M2D:4D, and D_[R-L]_) in females, and no such effects were observed in males.

The only other study that reports on an association between amniotic sex hormone levels and mothers’ 2D:4D^11^ found a negative correlation with T. The findings from our study generally contradict this observation, as the only significant correlations observed between amniotic T (specifically for LCMS measurements when the foetus was female) and maternal 2D:4D were in the opposite (i.e. positive) direction. These effects should of course be interpreted with considerable caution: not only are they in the opposite direction to that predicted by theory, but the corresponding correlations observed for the larger sample for which RIA T measurements were available are in the negative direction and not statistically significant. It should also be noted that relatively little consideration has yet been given to the possibility of associations between maternal 2D:4D and amniotic hormone concentrations.

The most notable finding from the current study is that neither T nor E, nor the T:E ratio present in amniotic fluid was a significant predictor of children’s 2D:4D ratios measured at 4½ year follow-up. This observation may be interpreted in several ways. Firstly, it could be that prenatal sex hormone exposure does indeed influence the development of 2D:4D, but that such processes occur earlier in pregnancy (i.e. towards the end of the first trimester)^17^. Support for this idea comes from the finding that 2D:4D already shows sexual dimorphism by the 9^th^-12^th^ week of gestation^18^. However, as there appears to be a certain amount of lability in 2D:4D during infancy^19^ and childhood^20^, postnatal exposure to sex hormones may also play a role. A second possible explanation for the current null findings is that second trimester sex hormones do influence the development of 2D:4D but that the concentrations measured in amniotic fluid simply do not accurately index those present in the foetal circulation^21^. A third possibility is that prenatal T and E do not determine variation in 2D:4D ratios (or that any association between these variables is smaller than initially thought).

It is noteworthy that we found no association between amniotic sex hormone concentrations and the children’s right-left difference in 2D:4D (D_[R-L]_). Although initially suggested by Manning^11^ as a further indicator of prenatal androgen action nearly two decades ago, there has been relatively little research into the validity of this proposed marker. Although we did find some evidence for D_[R-L]_ being lower in males than females, our findings are consistent with previous studies showing that this measure does not correlate with T measured from amniotic fluid^4^, maternal circulation^4,22^ or umbilical cord blood^22,23^, and that it does not differ between patients with congenital adrenal hyperplasia and unaffected controls^5^. As D_[R-L]_ is calculated as a ratio from two other noisy markers, its reliability can also be problematic^24^. Taken together, these observations raise serious questions regarding the utility of D_[R-L]_ as an indicator of prenatal androgen exposure.

The current findings should be considered in light of some important limitations. Firstly, the method used for measuring finger lengths in this study was unusual in that some participants were measured directly, others were measured indirectly (i.e. from photocopies), and a subsample was measured using both techniques. It is therefore important to note that digit ratios measured from photocopies are typically lower (i.e. more male-typical) than those measured directly^25^. However, we corrected for this problem mathematically, and so all participants could be examined simultaneously. It should also be noted that T analyses specific to the direct and indirect measurements of finger lengths from this cohort have been reported in an unpublished MPhil thesis^16^, and showed the same pattern of (null) results as reported here. Another limitation was that we could only examine E and T:E concentrations in a subsample, meaning that the statistical power for the associated analyses was lower than that for the RIA T analyses. However, it should be noted that the only other study to report associations between amniotic E (and T:E)^13^ had a very similar sample size (n=29; current sample for E/T:E: n=30).

## 5. Summary

The current study attempted to replicate the finding of Lutchmaya et al.^13^ that T:E ratio in mid-trimester amniotic fluid was a significant negative correlate of children’s 2D:4D ratios. However, we found no evidence that individual differences in amniotic T, E, or T:E ratio could predict children’s digit ratios measured at 4½ years of age. We did observe some correlations between amniotic T (and T:E ratio) and maternal digit ratios, though the direction of these effects was more often than not in the opposite direction to that which would be predicted by theory. Furthermore, we observed no correlation between amniotic sex hormones and the children’s right-left difference in 2D:4D (D_[R-L]_). Taken together, the findings suggest that mid-trimester amniotic T and E do not significantly influence development of the 2D:4D ratio. However, the possibility remains that these hormones do influence the development of digit ratio at an earlier stage of gestation.

## Acknowledgements

The authors wish to extend their gratitude to Prof Melissa Hines and Prof Vivette Glover for providing us with access to these data; we would also like to thank all of the parents and children who took part in this research.

## Notes

### Competing Interest Statement

The authors have declared no competing interest.

